# Protein features identification for machine learning-based prediction of protein-protein interactions

**DOI:** 10.1101/137257

**Authors:** Khalid Raza

## Abstract

The long awaited challenge of post-genomic era and systems biology research is computational prediction of protein-protein interactions (PPIs) that ultimately lead to protein functions prediction. The important research questions is how protein complexes with known sequence and structure be used to identify and classify protein binding sites, and how to infer knowledge from these classification such as predicting PPIs of proteins with unknown sequence and structure. Several machine learning techniques have been applied for the prediction of PPIs, but the accuracy of their prediction wholly depends on the number of features being used for training. In this paper, we have performed a survey of protein features used for the prediction of PPIs. The open research challenges and opportunities in the area have also been discussed.

## 1 Introduction

Proteins are important for functioning of our body. The structure of a protein influences its function by determining the other molecules with which it can interact. The protein interactions can reveal hints about the function of a protein. Protein-protein interactions (PPI) play an important role in living cells that control most of the biological processes, and most essential cellular processes are mediated by these kinds of interactions. Proteins mostly perform their functions with the help of interactions with other proteins. For example, disease-causing mutations that affect protein interactions may lead to disruptions in protein-DNA interactions, misfolding of proteins, new undesired interactions, or enable pathogen-host protein interactions. Similarly, aberrant protein-protein interactions have caused several neurological disorders including Parkinson and Alzheimer’s disease. With appropriate knowledge of interaction, scientist can easily predict pathways in the cell, potential novel therapeutic target, and protein functions. Hence, these examples have motivated to map interactions on the proteome-wide scale. The prediction of PPI has emerged as an important research problem in the field of bioinformatics and systems biology.

A PPI network focuses on tracing the dynamic interactions among proteins, thereby illuminating their local and global functional relationships (Rao et al., 2014). Experimentally determined PPI network from high-throughput techniques, such as yeast two hybrid (Y2H) screens and Tandem affinity purification coupled to mass spectrometry (TAP-MS), are inherently noisy that contains a large number of false positives. Predicting PPIs using experimental techniques are time-consuming, costly, need man-power, and also unreliable. Therefore, computational methods for the prediction of PPIs have evolved. Among computational techniques, machine learning has been extensively used for several classification and prediction problems, including PPIs. Machine learning is a data-driven approach which requires sufficient number of training sets and features. It has been found that number of protein features play vital role in the accuracy of prediction algorithms. Therefore, it is required that we must identify various protein features that need to be used to train a machine learning algorithm.

This review presents 13 different protein features which can be used for the prediction of PPIs. The paper is organized as follows. Section 1.1 and 1.2 describes types of PPIs and experimental methods for finding PPIs, respectively. Section 2 presents a brief description of computational approaches for the prediction PPIs. Section 3 covers 13 different proteins features that can be used for training machine learning algorithms. Section 4 describes how protein features are represented so that it can be fed to a learning algorithm. Finally, section 5 concludes the paper and discusses challenges and opportunities in the area.

### 1.1 Types of protein–protein interaction

It is analyzed and found that determination of PPI by different methods show a low degree of overlap. Hence, researchers have designed computational tools to assess the reliability of data coming out of high-throughput experiments. This low overlapping of interaction data and low reliability of high-throughput experimental techniques show that PPI determined with various approaches explore different types of interactions. De Las Rivas and de Luis (De Las Rivas & de Luis, 2004) classified the PPI into three level of association with several sublevels, as shown in Fig. 1.

**Fig. 1.**
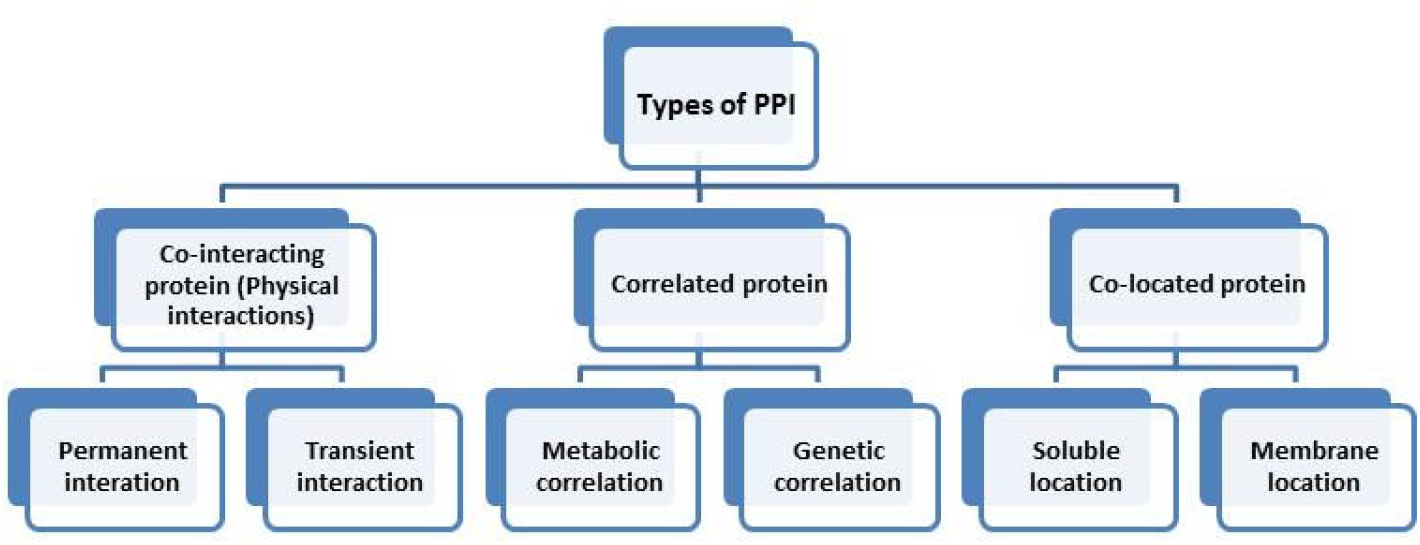
Classification of protein-protein interaction (Source: De Las Rivas & de Luis, 2004)

*Physical interactions* are that where proteins form a stable protein complex and performs biomolecular role such as structural and functional role. They are protein subunits of the complex that work together. *Correlated proteins interactions* are those which do not interact physically but are involved in the same biomolecular activities. Correlated protein interactions may be metabolic correlation or genetic correlation. *Co-located proteins interactions* are those where proteins are defined to work in the same cellular compartment.

### 1.2 Experimental methods

There are two main technologies that determine PPI interactions: binary method and co-complex method. Binary techniques measure direct physical interactions between protein pairs. Yeast Two Hybrid (Y2H) method is one of the mostly used binary method. Co-complex method measures physical interaction among groups of proteins, without finding pairwise interactions. Tandem affinity purification coupled to mass spectrometry (TAP-MS) is most often used Co-Complex method. Generally, Co-complex methods measures both direct and indirect interactions between proteins. The experimental result of both the methods is totally different from each other.

Data obtained from co-complex method cannot be directly mapped to binary interpretation. An algorithm or model is required to map group-based data into pairwise interactions. The ‘spoke model’ is most widely applied technique to transform data from group-based to pairwise interactions (de Las Rivas & Fontanillo, 2010).

## 2 Computational Predictions

Mostly proteins do their functions by interacting with other proteins. The PPI within a cell may enrich our understanding about protein functions and cellular processes. Over the past few years, due to advancement in computation biology and bioinformatics, an explosion in functional biological data derived from high-throughput technologies to infer PPI has been observed.

Many large-scale experimental techniques have been employed to study PPIs including Y2H screens, X-ray crystallography, NMR and site-directed mutagenesis. But these experimental techniques are costly, tedious, time-consuming, labor-intensive and potentially inaccurate (Browne et al, 2006; Wang et al, 2013). On the other hand, tremendous protein interaction data has been generated out of proteomics research that need to be validated and annotated structural information.

Computational methods play a significant role in the prediction of PPI. They are used to predict potential interactions between proteins, to validate results of high-throughput interaction screens and to analyze the protein networks inferred from interaction databases. Several statistical and machine learning based methods have been applied for the prediction of PPI including Bayesian Networks (Jansen et al., 2003; Patil & Nakamura, 2005), Simple Naïve Bayesian, Random Forest (Šikic et al. 2009; Zubek et al., 2015), Support Vector Machine (Bock & Gough, 2001; Chatterjee et al., 2011; You et al., 2013; You et al., 2014; Zubek et al., 2015), Decision Tree, Logistic Regression, k-Nearest Neighborhood (kNN), Conditional Random Field, Artificial Neural Networks (Fariselli et al., 2002), to name a few. Despite the success of these methods, there is still need for the improvement in terms of prediction accuracy and computational efficiency (Res et al, 2005, Bordner & Abagyan, 2005).

PPI predictions can be accomplished broadly in four steps, as shown in Fig. **2**. Initially protein features will be extracted from different genomic and proteomic information. These protein features are represented in the form of a vector so that it can be used to train a machine learning classifier. Once a classifier is properly trained with the extracted protein features, it must be compared with the gold standard to assess its performance. A detail description of data-driven approach for prediction of PPIs can be found in Xue, et al. (2015).

**Fig. 2.**
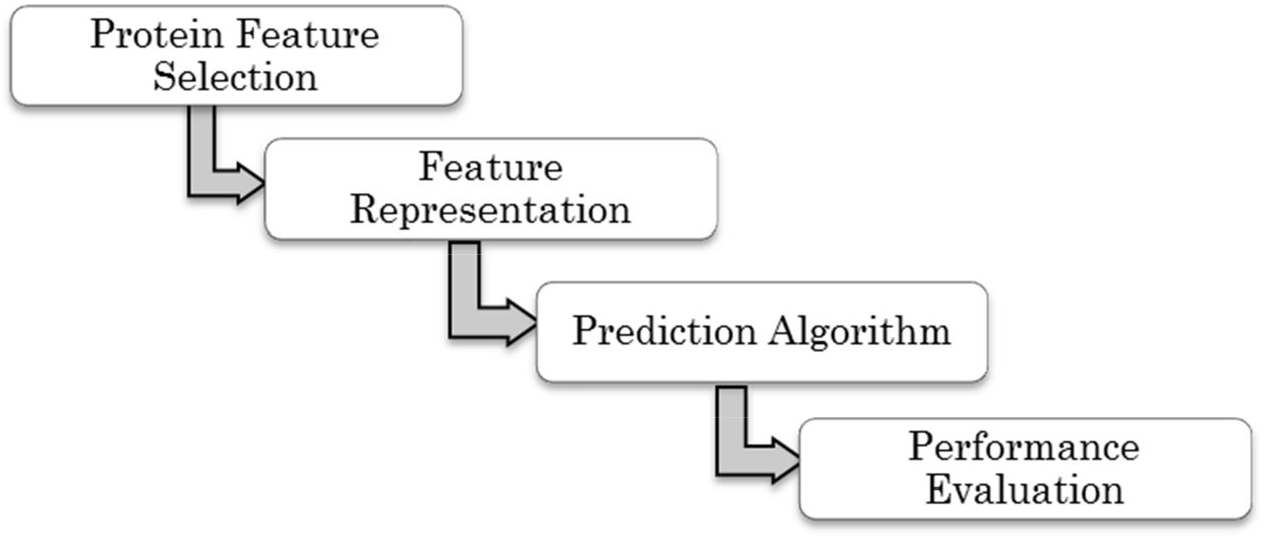
Generic steps in machine learning-based protein-protein interaction predictions

## 3 Protein Features Selection

In the literature, various protein features have been used to predict PPI, either individually or in combinations. It has been found that none of the single protein feature is sufficient to predict PPI because single feature alone does not carry adequate information. Hence, a combination of some of these features has been found to be a better way to enhance the performance of machine learning for PPI predictions (Wang et al, 2006; Browne et al, 2006; Sikic et al, 2009). Detail of the protein features are described as follows:

### 3.1 Primary sequence

The primary sequence of protein corresponds to linear amino acid sequence. In the literature, initially PPI prediction has been carried out by using sequence information only (Ofran & Rost, 2003; Sikic et al., 2009). Ofran & Rost (2003) proposed a neural network based approach to identify PPI interfaces from sequence information. The result shows 94% accuracy for most strongly predicted sites. The results of Ofran & Rost (2003) indicated that PPI sites are possible to be predicted using sequence alone. Sikic and colleagues (Sikic et al., 2009) have used names of nine consecutive residues in a sequence as input feature vector. The input feature vector was defined on a sliding window of 9 residues. The window was considered as positive, if at least N residues including the central residue were marked as interacting. The data was classified for values of N ranging from 1 to 9 and result was evaluated. The input vector consists of all nine residues’ names, and min, max or average value of features. When Random Forest classifier has been applied, the result shows precision of 84% and recall of around 26%. However, accuracy of the method is not good. In this direction, researchers suggested that we cannot predict PPI with acceptable accuracy using sequence information only. We need to incorporate other information, such as evolutionary and structure information, to predict PPI with high accuracy.

### 3.2 Secondary and 3D structure

Proteins are polymers comprising of chains of amino acids. The structure of proteins determines their specific function. The structure of the protein can be described at different levels. The first level is the primary structure that corresponds to linear amino acid sequence. At the second level, we have secondary structure that refers to how the amino acid back-bone of the protein is arranged in 3-dimensional space, by forming hydrogen bonds with itself. There are three common secondary structures: alpha helices, beta sheets and random coils. Third level is tertiary structure which is produced when elements of the secondary structure fold up among them. Finally, the quaternary structure is related to the spatial arrangement of several proteins.

Sikic and his colleagues (Sikic et al., 2009) combined sequence information and 3D structure information to predict PPI using Random Forest classifier. They have used all 3D structure information available from Protein Structure and Interaction Analyzer (PSAIA) developed by Mihel and colleagues (Mihel et al., 2008), in additional to secondary structure. A total of 26 features have been considered here for training the classifier. Since, Random Forest algorithm is capable to estimate the importance of a particular feature, so they have applied input parameter set reduction. Here, it has been noticed that information acquired from sequence has the highest importance in the prediction. Also, five best structure features has been ranked as: non-polar accessible surface area (ASA), maximum depth index, relative non-polar ASA, average depth index, and minimum protrusion index.

### 3.3 Sequence entropy

Variability within related protein sequences has been proved to provide clue about their 3D structure and function. The high variability regions within related proteins groups are linked to the specificity of molecules, while low variability regions are mostly structural or define regions of common function. Shanon entropy can be applied to estimate the diversity among group of protein sequences. For alignment of multiple sequences, the entropy *H* for each position can be computed as,

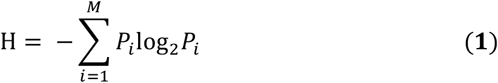

where, *P*_*i*_ is fraction of residues of amino acid type *i*, and *M* is total number of amino acid type, i.e., 20. The sequence entropy score basically ranks the frequencies of the occurrence of 20 different types of amino acid, where lowest values correspond to the most conserved positions. The HSSP (Homology-derived Structures of Proteins) is a derived database that merges 3D structure and sequence information. This database is a noble resource for extracting sequence entropy (Schneider & Sander, 1996).

### 3.4 Evolutionary conservation

Conserved sequences are similar or identical sequences that may have been maintained by evolution despite speciation, if it is cross species conservation. More highly conserved sequence may occur in the phylogenetic tree. Since sequence information is normally transmitted from parents to progeny by genes, a conserved sequence implies that there is a conserved gene. There are several methods that use evolutionary conservation as a primary indicator to find location of interface residue in PPI. The main reason for using evolutionary conservation is that it reflects evolutionary selection at the interaction sites to maintain functions of protein families (Lichtarge et al., 1996; Neuvirth et al., 2004; Wang et al., 2013). At the interfaces, evolutionary conservation of residue is observed as higher than general surface residues, and hence has distinct feature of protein interaction sites (Wang et al., 2013).

### 3.5 Solvent accessible surface area

The accessible surface area (ASA), also known as solvent-accessible surface area (SASA), is the surface area of a biomolecule which is accessible to a solvent (Connolly, 1983; Richmond, 1984). Measurement of SASA is mostly measured in terms of square angstroms and was initially explained by Lee and Richards in 1971 (Lee & Richard, 1971). SASA is computed using the 'rolling ball' algorithm proposed by Shrake & Rupley (1973). This algorithm applies a sphere of a particular radius to 'probe' the surface of the molecule. Some of the other methods to calculate SASA are linear combination of pair-wise overlap (LCPO) (Weiser et al., 1999) and Power Diagram method. The relative SASA may be applied to estimate the magnitude of binding-induced conformational changes using structures of either monomeric proteins or bound subunits. When it is applied to a large set of complexes, the result shows large conformational changes due to common binding. The SASA including many other protein features have been used in Sikic et al., 2009 for finding PPI sites.

### 3.6 Protein expression

The co-expression of genes may act as an indicator of functional linkage. Hence, Microarray mRNA expression data can be used to predict PPIs. Here, interactions between two proteins are predicted based on similar expression of their coding genes in multiple conditions. There are several methods using which we can compute the similarity among expression profiles of genes in the given gene expression matrix. Pearson’s Correlation Coefficient is a popular method to compute similarity since it can distinguish positively and negatively regulated gene pairs. The idea here is to identify set of genes with similar expression patterns.

This microarray mRNA co-expression dataset is based on two assumptions: i) proteins present in same complex usually interact to each other, and ii) proteins belong to same complex are co-expressed. One of these datasets has been reported by Cho et al. (1998). The dataset represents time-course gene expression fluctuations during yeast cell cycle and Rosetta compendium, comprises of expression profiles of 300 deletion mutants and cells under various chemical treatments (Cho et al, 1998). Cells have been collected at 17 time points observed at 10 min intervals, covering almost two full cell cycles. Here, pair-wise Pearson’s correlation were computed for every gene-pairs, value of which ranges between 0 and 1.

### 3.7 Marginal essentiality

At the most basic level, functional significance of genes may be defined by its essentiality, i.e., genes which are knocked out may render the cell unviable. Marginal essentiality is basically based on a hypothesis, called ‘marginal benefit’, that several non-essential genes make small but significant contributions to fitness of a cell (Browne et al., 2006). Marginal essentiality ‘M’ can be defined as a quantitative measure of a non-essential gene’s importance to a cell (Yu et al., 2004). Yu et al. (2004) described that marginal essentiality measure relates to several topological properties of PPI networks. Particularly, proteins having higher degree of *M* tend to be hubs of the network, and have a shorter characteristic path length to their neighbors. Yu et al. (2004) obtained marginal essentiality dataset by combining the result of four phenotypic experiments. Two proteins are considered to be interacting, if they have higher combined *M*.

### 3.8 Co-essentiality

This feature is based on the hypothesis that proteins are either essential or non-essential, indicating that they belong to the same complex. If this is the case, they may be either essential or non-essential but not both. The reason is that a deletion mutant of either protein would yield same phenotype, and their mutual deletion would harm the function. In Zhang et al. (2012), co-essentiality, along with several proteins features, has been considered for the prediction as well as analysis of Protein Interactome in Pseudomonas aeruginosa. Gene essentiality data were retrieved from Database of Essential Genes (Zhang & Lin, 2009). Each gene of PA PAO1 (a pathogen P. aeruginosa which is problematic in chronic airway infections) is considered as either essential (678 genes) or non-essential (4,890 genes). Considering gene essentiality, each gene-pair has a nominal value for co-essentiality: both are essential, both are non-essential, one is essential and other is non-essential.

Browne et al. (2006) has derived co-essentiality dataset from both MIPS complex catalogue and transposon and gene deletion experiments. In this dataset, protein-pairs are assumed to interaction, if both proteins are either essential or non-essential. When there is a combination of essential and non-essential proteins, protein-pair is considered as non-interacting. In Browne et al. (2006), mixture of non-essential and essential protein-pairs is represented by 1 and only non-essential are presented by 0. More detail information about MIPS based co-essentiality data can be found in Mews et al. (2000).

### 3.9 MIPS functional catalogue

It is a hypothesis that protein-pairs acting in same biological processes are more likely to interact to each other. Hence, based on this hypothesis, protein-pairs can be defined as interacting, if they belong to same biological process. Functional similarity between two proteins has been estimated from Gene Ontology (GO) and MIPS functional catalogue (FunCat). MIPS FunCat is annotation scheme that represents functional description of proteins and contains 28 main functional categories covering cellular transport, metabolism and signal transduction. In other words, FunCat is a hierarchically organized, organism independent, flexible, scalable and controlled classification approach that enables functional description of proteins used for manual annotation of prokaryotes, eukaryotes, animals, fungi and plants. The main categories are organized as a hierarchical tree-like structure that describes up to 6 levels of increasing specificity. The FunCat version 2.1 includes a total of 1362 functional categories. Here, each protein represents a subtree of overall hierarchical class-tree. Now, it is possible to compute intersection tree of two subtree for two given proteins. This estimation is similarly done for complete list of protein pairs. For example, there are ∽18 million interactions in yeast. The FunCat is separate from MIPS complex categories. The functional similarity between protein-pairs can be finally described as frequency at which interaction trees of protein-pair occur in the distribution.

### 3.10 GO-driven frequency based similarity

By consider the hypothesis, as considered in FunCat, that is, protein-pair involved in same biological process are likely to interact; it is possible to extract participation information of protein-pairs in specific biological process using Go-driven annotation databases such as Saccharomyces Genome Database (Cherry et al., 1998). Browne et al. (2006) applied GO-driven Frequency Based Similarity measures to generate dataset to predict PPIs. This dataset was obtained by computing the similarities between gene products annotated in GO biological process hierarchy. Consider two proteins *p*_*1*_ and *p*_*2*_ which has a specific set of lowest common ancestor nodes in hierarchy, then it is possible to count number of proteins pairs having same set of annotation terms. In Browne et al. (2006), for all possible protein pairs in *S. Cerevisiac* (∽18 million), they counted how many of these possible pairs share exactly same functional terms resulting a number ranges between 0 and 18 million. A smaller number represents a more specific functional description of a protein pair that suggests higher functional similarity and higher possibility to belong to same complex. On the other hand, a larger value of count represents a less functional similarity and there is less possibility for the protein-pair to belong to same complex.

### 3.11 GO-driven semantic similarity

To calculate similarity between Go terms and gene products, Lin’s semantic similarity technique (Lin, 1998) is mostly applied (Azuaje et al., 2005). The similarity values of gene-pairs can be used for PPI prediction, as used in Browne et al. (2006), to construct GO-driven Semantic Similarity (GOSEM) dataset. The GOSEM method exploits information contents of shared GO terms parents, as well as that of query GO terms applied in annotating gene. This similarity measure is expressed as probability and based on count of each terms occurring in the GO corpus. Lin’s method computes similarity between two GO terms *x*_*i*_ and *x*_*j*_ as,

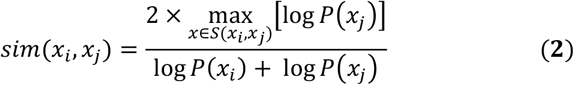

where *S* is set of parental terms shared by terms *x*_*i*_ and *x*_*j*_; *P*(*x*) is probability of finding terms *x* or any of its parent in the GO corpus. The similarity between genes can be computed by aggregating similarity values of annotated terms of genes. The similarity between gene products *g*_*k*_ and *g*_*l*_ can be defined as average interest similarity between terms *A*_i_ to *A*_*j*_,

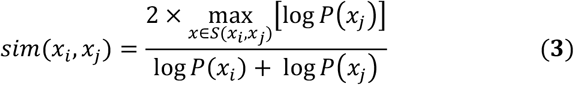

where, *A*_*k*_ and *A*_*l*_ comprises of *m* and *n* terms.

### 3.12 Position-specific scoring matrices

Position Specific Scoring Matrices (PSSMs), also called Position-Specific Weight Matrix (PSWM), are most commonly applied motifs descriptor in biological sequences. It is often calculated from a set of functionally related and aligned sequences. PSSM attempts to capture intrinsic variability characteristic from multiple sequence alignment (MSA). PSSMs have been used as protein feature vector by many researchers for finding PPIs (Deng et al., 2009). A PSSM has one row for each residue (4 rows for nucleotides or 20 rows for amino acids). Given a set *S* (*S*={*s*_*1*_, *s*_*2*_, …, *s*_*n*_}) of *n* aligned sequences having length *l*, PSSM can be calculated as,

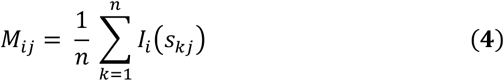

where, *i*=A, C, T, G or amino acid residues; *j*=1, 2, …, *l*; *s*_*k*_=*s*_*k1*_, *s*_*k2*_, …, *s*_*kl*_ (*s*_*ki*_ being any of the residue). Each coefficient in PSSM indicates count of a residue at a given position.

### 3.13 Residue interface propensity

Residue Interface Propensity (RIP) quantifies preferences of an amino acid to act as interface site of two interacting proteins. It is calculated by using interaction of all the proteins in entire family. For each amino acid *i*, the propensity is defined as,

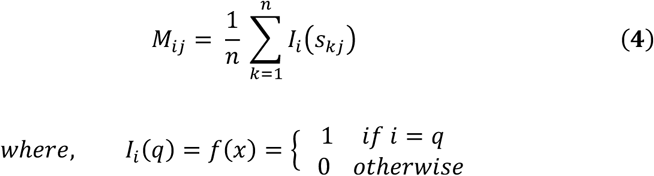

where *N*_in_(*i*) is number of amino acid of type *i* in interface; *N*_*in*_ is total number of amino acids of any type in the family; *N*_surf_(*i*) is number of surface amino acid of type *i* in all domain; and *N*_surf_ is total number of amino acids of any type in the family. In general, *Propensity*_*i*_ > 1.0 indicates that residue *i* has high chance for being in interface. The propensity score depends on both, number of interaction inferred from PSIMAP and size of SCOP family. RIP has been applied by Dong et al. (2007) and Lui et al. (2009) to identify protein binding sites.

## 4 Protein Feature Representations

There are different proteins features, such as sequence information, co-expression, protein structure information, phylogenetic profiles, and so on, that can be used to predict PPIs computationally. Once protein features has been selected, the next fundamental question is how these protein features are represented so that it can be fed to a classifier for training and prediction. For the representation of sequence information as feature vector, a sliding window is most deployed approach to represent association among neighboring residues. Selecting length of sliding window is a vital issue because it affects the prediction accuracy. There is no thumb rule for the selection of length of sliding window and mostly it is set randomly. In a recently study, Sikic et al., 2009 applied an entropy based method to find out window length, which is given by,

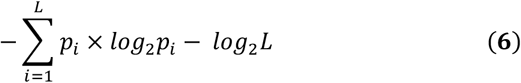

where, *L* is window length, and *p*_*i*_ represent the appearance frequency of *i*_*th*_ interacting residues in window of *L* residue given a central interaction. Results observed by Sikic et al. (2009) was similar for different window length but window length of 9 shows maximum difference of entropy. You et al. (2014) proposed a novel feature representation technique. It has been assumed here that continuous and discontinuous amino acids segment play a vital role in predicting PPI. The proposed feature representation approach considers interaction between sequentially distant but spatially closed residues. Here, Multiscale Continuous and Discontinuous (MCD) method is deployed for sequence representation to transform protein sequence to feature vector using binary encoding scheme. A multiscale decomposition technique is applied to divide given sequence into chunk of multiple sequences of distinct length. For the extraction of information, entire protein sequence is divided into equal length segments, and further a binary coding scheme is applied to each segment. Each segment is encoded as 4-bit binary sequence of 1’s or 0’s. For every continuous or discontinuous region, composition (C), transition (T) and distribution (D) – three kinds of descriptors are applied to represent its characteristics. Here, ‘C’ is ratio of number of amino acids in local region. ‘T’ specifies frequency (in percent) to which amino acids of a particular property is followed by that other property. ‘D’ means the length of chain within first 25%, 50%, 75% and 100% of amino acids of particular property are located.

## 5 Conclusion and Discussion

It has been observed that there are still discrepancies in estimating the number of PPIs even for small size well-studied unicellular organism Saccharomyces cerevisiae (yeast). Different studies estimated different size of complete binary protein interactome of yeast containing ∽6,000 proteins. Grigoriev (2003) estimated around 16,000 to 26,000 PPIs, on the other hand Blow (2009) reported more than 30,000 potential interactions in yeast. Some of the databases have only experimental data containing greater than 50,000 binary PPIs in yeast. Thus, this discrepancy in the results indicates that some of the experimentally obtained interactions most probably contain false positives. Machine learning algorithms are playing a pivotal role in several classification and prediction problems including PPIs predictions. Prediction of PPI is a non-trivial problem because there are lots of factors which are involved in binding of amino-acids. Feature selection is an important issue in machine learning techniques, specially protein features in PPI prediction problem. This review covers a maximum of 13 different protein features used in the literation.

In case of incomplete datasets, careful study is required to know the optimal machine learning technique, and optimal protein features. Some of the suggested future directions are as follows:

i. Most of the methods proposed earlier consider either single or few protein features for training classifiers. The results show that considering multiple protein features improves the performance of PPI prediction. Hence, we can look forward for integrating more protein features for further improvement in the prediction results.
ii. Today, several ensembles learning algorithms have been developed, including boosting, bagging and multi-classifier systems, showing better prediction accuracy in various classification and prediction problems. Hence, we can deploy these algorithms to increase the prediction accuracy at the cost of computation.
iii. Mostly of the studies have been performed on finding PPIs on yeast only. We can look for finding PPIs in *Homo sapiens* in similar way.

## Acknowledgments

This work is financially supported by Jamia Millia Islamia, New Delhi, India under innovative research activities.

